# Translational Fidelity Maintained by ZNF598-Dependent Ribosome Resolution Enables Cellular Adaptation to High Energy Demand

**DOI:** 10.64898/2025.12.19.695646

**Authors:** Toufic Kassouf, Laura Ryder, Katia El Ghoz, Bartosz Artur Kucharski, Lucile Dollet, Filip Dziaczkowski, Goda Snieckute, Sijie Chen, José Francisco Martínez, Dominika Malińska, Angel Loza-Valdes, Malgorzata Alicja Sliwinska, Eric Bennett, Zach Gerhart-Hines, Simon Bekker-Jensen, Grzegorz Sumara

**Affiliations:** Dioscuri center for metabolic disease, Nencki Institute of Experimental Biology, Polish Academy of Sciences, Warsaw, Poland; Center for Gene Expression, Department of Cellular and Molecular Medicine, University of Copenhagen, Blegdamsvej 3, DK-2200 Copenhagen, Denmark; School of Biological Sciences, Department of Cell and Developmental Biology, University of California, San Diego, La Jolla, California, USA; Novo Nordisk Foundation Center for Basic Metabolic Research, Faculty of Health and Medical Sciences, University of Copenhagen, Blegdamsvej 3B, DK-2200 Copenhagen, Denmark

**Author notes:** Correspondence should be addressed to Grzegorz Sumara and Simon Bekker-Jensen.

## Abstract

Cellular adaptation to stress evoked by high energy demand requires tight coordination between transcriptional programs, protein synthesis, and organelle function. In many physiological contexts, metabolic remodeling depends on increased mitochondrial capacity and activity, placing high demands on the translational machinery. Beige and brown adipocytes, which undergo rapid mitochondrial expansion and activation in response to physiological cues, provide a powerful model to study how cells adapt their translational output to meet increased metabolic demand. How translational fidelity is maintained, and stress-associated conflicts in protein synthesis are resolved during such adaptive processes, remains incompletely understood.

Here, we identify ZNF598 as a key factor that resolves translation stress caused by ribosome collisions during metabolic adaptation of adipocytes. We show that diverse physiological stimuli, including hormonal signaling, environmental challenges, and dietary changes, induce ZNF598 and associated enzymes in metabolically active cells. ZNF598 is required for efficient mitochondrial biogenesis and function, supporting adaptive increases in respiration and energy expenditure. Loss of ZNF598 compromises these adaptive responses and leads to metabolic dysfunction, whereas enhancement of ZNF598 activity improves cellular and organismal metabolic flexibility. Together, these findings indicate that distal steps of the translational machinery are a rate-limiting factor for adaptation to increased energy demand.

## INTRODUCTION

Tissue remodeling requires the acquisition of new organelles, which is associated with the activation of transcriptional and translational programs. Transcriptional outputs are tightly regulated by a plethora of transcription factors and co-factors. However, the regulation of translational output, which is ultimately required for the production of new proteins, is less well understood. Adipose tissue represents a perfect example of a highly plastic organ that undergoes dynamic changes in response to different physiological states^1^. Adipose tissue is central to whole-body metabolism. Three main functional types of adipocytes are distinguished. White adipocytes store large quantities of energy in the form of triglycerides (TG) while beige and brown adipocytes additionally dissipate energy in the form of heat^2^. Thermal or dietary challenges can trigger various hormonal-induced signals, including β-adrenergic-dependent signaling, that drive the transition from white to beige adipocytes. In brown and beige adipocytes, β-adrenergic signaling boosts mitochondrial content and activity to sustain adaptive thermogenesis. Similarly, β-adrenergic signaling induces the expression of uncoupling protein 1 (UCP1), which disrupts the proton gradient used for ATP synthesis, resulting in energy dissipation as heat^2^.

While the induction of energy dissipation by adipocytes represents an attractive strategy to combat obesity and diabetes^2^, from an evolutionary point of view this process is likely subject to multiple layers of control mechanisms to prevent unnecessary nutrient waste. The first layer is exerted by factors that sense and transmit hormonal and physiological inputs to transcriptional effectors. Cold exposure and β-adrenergic stimuli induce, among dozens of other transcription factors, Peroxisome Proliferator-Activated Receptor-Gamma Coactivator-1 α (PGC1α)^3^ and PR domain containing 16 (PRDM16)^4^ to upregulate expression of mitochondrial genes and *UCP1* in adipocytes. The proteins encoded by these transcripts support oxidative phosphorylation and thermogenic capacity. While considerable attention has been given to transcriptional events occurring during adaptive thermogenesis, regulation at the translational level is much less studied. Consequently, the scale at which translation and ribosomal output is rewired upon hormonal and environmental cues in brown and beige adipocytes remain poorly understood. Previous studies indicate that exposure to cold or β-adrenergic stimulation enhances ribosomal output in adipocytes^5^ with unknown consequences for translation fidelity.

Cells have evolved a suite of surveillance pathways to detect and resolve translation conflicts. These include specific responses to stalled and collided ribosomes, which can result from ribosomal congestion on problematic mRNA sequences, under-availability of amino acids and tRNAs, oxidative stress, or genetic mutations^6–8^. Collided ribosomes are sensed by cells as a proxy for aberrant translation and trigger ribosome-associated quality control (RQC)^6,7,9,10^. The RQC machinery is conserved from yeast to human cells and is related not only to cytosolic protein quality control but also to mitochondrial function and protein aggregation^7,9^. The unique structure of collided ribosomes is recognized by the E3 ubiquitin ligase ZNF598 that promotes ubiquitination of the ribosomal proteins uS10 (RPS20) and eS10 (RPS10)^11–13^. ZNF598-ubiquitinated ribosomes accrue the RQC-trigger (RQT – ASCC2, ASCC3, and TRIP4^14–16^) complex for resolution, thus enabling trapped nascent chains to be extracted and degraded and ribosomal subunits to be recycled^6,7,9,10^. Separate from ZNF598-mediated ribosome ubiquitination, the E3 ligase RNF10 ubiquitinates uS3 (RPS3) and uS5 (RPS2) to promote degradation of 40S subunits engaged in non-productive mRNA binding (initiation RQC, iRQC). These may result from ZNF598-mediated resolution of collided ribosomes, impaired 40S initiation scanning, or prolonged nutrient starvation^14^. RNF10-mediated ubiquitination is counteracted by the deubiquitinating enzyme USP10^17,18^, and 40S decay requires RIOK3 engagement with ubiquitinated 40S^19–21^. Following resolution of the leading ribosome in the collision structure, the remaining 80S ribosome is dissociated by the ribosome recycling factors

Pelota, HBS1L, and ABCE1^6,7,9,10^. While 40S subunits can either be directly recycled for subsequent rounds of translation or degraded through RIOK3 action, 60S subunits remain associated with peptidyl-tRNA and nascent chains. These particles are recognized by the RQC component NEMF that binds tRNA exposed at the subunit interface of 60S-nascent chain complexes. This event facilitates binding of the E3 ubiquitin ligase Listerin (LTN1), leading to ubiquitination, p97-mediated extraction, and proteasomal degradation of the nascent chain^6,7,9,10^.

Apart from the severe and early-onset neurodegeneration associated with *Ltn1* and *Nemf* whole-body knockout in mice^22,23^, physiological contexts for ribosome collision and the importance of RQC for mammalian biology have yet to be appreciated. This includes its role in metabolism in general and in adipose tissue function, in particular. Our results highlight that adaptive thermogenesis is associated with a massive translational burst that is largely uncoupled from transcriptional gene activation. The resulting congestion of ribosomes on transcripts leads to widespread ribosome collision in adipocytes exposed to thermogenic stimulation. While RQC is dispensable for adipose tissue establishment during development, the pathway becomes critically important during adaptive responses such as white adipose tissue (WAT) browning and thermogenesis. We show that ZNF598 is required and sufficient for the induction of heat dissipation by mitochondria in brown/beige adipocytes. At the mechanistic level, ZNF598 supports the induced biogenesis of mitochondrial mass, mitochondrial proteins, and the uncoupler UCP1. Finally, conditional deletion of the *Znf598* gene in mouse adipocytes resulted in adipose tissue dysfunction and glucose intolerance, while overexpression of this factor prevented diet-induced diabetes and enhanced responses to β-adrenergic stimulation. In a broader physiological context, our data may suggest that energy-demanding cellular remodeling requires passage through a checkpoint that controls translational fidelity.

## RESULTS

### Stimulation of thermogenesis is associated with translational upregulation, ribosome collision, induction of ZNF598, and the RQC machinery in adipocytes

Activation of the adaptive thermogenesis program in brown and beige adipocytes by cold challenge, adrenergic stimulation, or dietary interventions requires remodeling of their subcellular structure and metabolism, incl. the acquisition of mitochondrial mass^3^. Stimulation of murine stromal vascular cells (SVC)-derived brown adipocytes, T37i-derived murine beige/brown adipocytes, and SVC-derived white adipocytes (Fig. 1A) with the β-adrenergic agonist CL-316,243 (CL) increased ribosomal output as evidenced by incorporation of puromycin into nascent chains (Fig. 1B; Fig. S1A,B). Correlating published transcriptome and proteome datasets from BAT tissue of mice exposed to acute cold (6 hours and 8 hours, respectively), we noticed that translational induction of mitochondria-related mRNAs appeared to dominate over transcriptional changes at this early timepoint (Fig. 1C). Such upregulated translational efficiency may compromise translational fidelity and engage the RQC pathway. Cold challenge of mice for 24 hours was accompanied by transcriptional up-regulation of *Znf598* (also referred to as *Zfp598*) and most other genes encoding components of the core RQC pathway, in both BAT and inguinal white adipose tissue (iWAT) (Fig. 1D,E). Moreover, the genes encoding the three known components of iRQC (RNF10, USP10, and RIOK3) were also significantly upregulated in BAT and iWAT by cold challenge (Fig. S1C,D). Of note, expression of genes related to the ribotoxic stress and integrated stress responses (RSR and ISR) ^24^ were not affected during the acute phase of the cold response in BAT and iWAT (Fig. S1C,D). This was except for the genes encoding the GCN1-associated ubiquitin ligases RNF25 and RNF14, which are also involved in quality control, specifically turnover of ribosomes with

**Figure 1.**
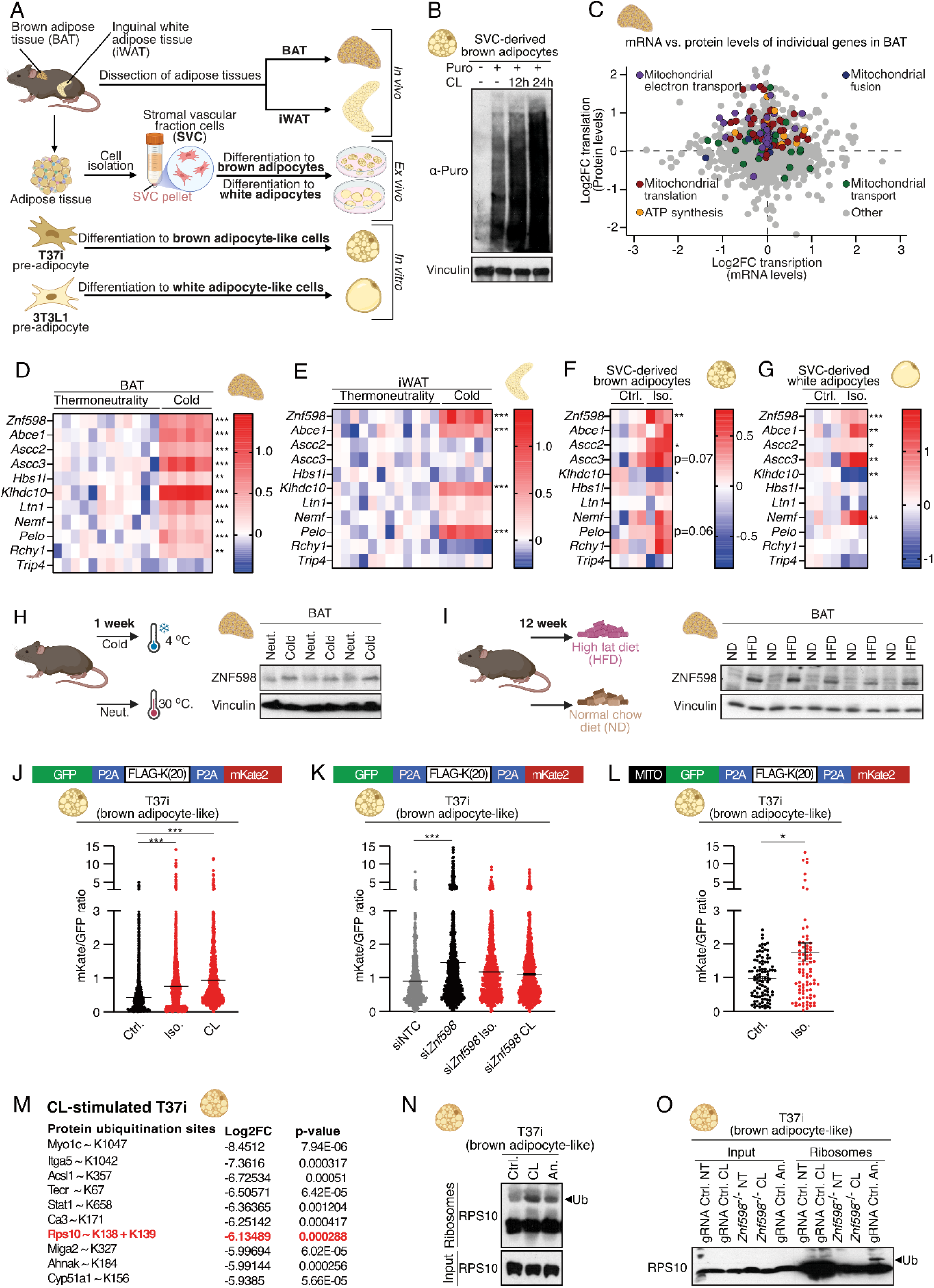
Translational Activation and Ribosome Quality Control in Thermogenic Adipocytes. **(A)** Experimental sources of mouse adipose tissue and adipocytes for this study. Whole brown (BAT) and white (WAT) adipose tissues were dissected from mice, stromal vascular cells (SVC) were isolated from mouse adipose tissue (white or brown) and differentiated into either brown or white adipocytes, and immortalized T37i and 3T3L1 pre-adipocyte cell lines were differentiated into brown and white adipocyte-like cells, respectively. **(B)** Determination of ribosomal output using puromycin-labeling of proteins in brown adipocytes derived from SVCs treated with CL-316,243 (CL) for the indicated time. **(C)** Correlation plot for mRNA vs. protein levels of individual genes in BAT of mice exposed for 6 (mRNA) and 8 (protein) hours to 4 °C. Genes and corresponding proteins associated with the indicated processes are shown with the following colors: mitochondrial electron transport chain (GO:0022904, “respiratory electron transport chain”) – purple, mitochondrial fusion (GO:0008053) – blue, mitochondrial translation (GO:0032543) – red, ATP synthesis (GO:0015986, “proton motive force-driven ATP synthesis”) – orange, mitochondrial transport (GO:1990542, “mitochondrial transmembrane transport”) – green, others – gray. **(D-G)** Heat maps showing the log2-transformed fold changes of *Znf598* mRNA levels and mRNA levels for other components of the RQC machinery in BAT **(D)** and iWAT **(E)** of mice kept at room temperature or exposed to 4 °C (n = 12 and n = 6 biological replicates, respectively) for 24 hours, as well as in SVC-derived brown **(F)** and white **(G)** adipocytes following β-adrenergic stimulation (isoproterenol – Iso.) for 6 hours (n = 3-4 biological replicates). **(H)** Western blot (WB) analysis of the abundance of ZNF598 in BAT collected from wild-type (WT) mice following 1 week of thermoneutrality (30 °C, neut.) or cold exposure (4 °C, cold). **(I)** WB analysis of the abundance of ZNF598 in BAT of WT mice fed a normal chow diet (ND) or a high-fat diet (HFD) for 12 weeks. **(J) and (K)** Top: schematic of a dual-fluorescent GFP-20K-mKate2 construct. GFP and mKate2 cDNA is split by a ribosomal stalling sequence containing 20 consecutive lysine (AAA) codons preceded by FLAG-tag cDNA. Bottom: Quantification of GFP to mKate2 ratio in BAT-differentiated T37i cells stably expressing the reporter and transfected with the indicated siRNAs (NTC, non-targeting control) and treated with Iso. and CL for 24 hours. Value from individual aggregates (GFP and/or mKate) from at least 10 randomly selected fields (5-8 cells per field) of view per sample are plotted, and lines indicate the mean values +/- standard error of the mean (SEM). **(L)** Top: Schematic of a dual-fluorescent GFP-20K-mKate2 stalling reporter with N-terminal mitochondrial targeting signal (MITO). Bottom: Quantification of GFP to mKate2 ratio in BAT-differentiated T37i cells stably expressing the reporter and treated with Iso. for 24 hours. Values from individual cells are plotted, and lines indicate the mean values +/- SEM. **(M)** Ubiquitination sites most affected by CL-treatment (24 hours – mock divided by CL, n = 3 biological replicates) of T37i brown adipocytes knockout (KO) for *Znf598* (enrichment of diGly remnant on peptides after trypsin digestion). The ZNF598-dependent and RQC-related ubiquitination sites on RPS10 are highlighted in red. **(N and O)** Western blot (WB) analysis of protein lysates (input) and pelleted ribosomes from T37i brown adipocytes WT (gRNA Ctrl.) or KO (^-/-^) for *Znf598*. Differentiated cells were treated with CL (24 hours) or anisomycin (An., 1 hour) as a positive control. (D-G, M) statistical analysis was performed using DESeq2 (v. 1.34), *, p≤0.05; **, p≤0.01; ***, p≤0.001 in Wald test. (J-L) *, p≤0.05; **, p≤0.01; ***, p≤0.001 in Mann-Whitney test or student’s t-test.

EEF1A trapped in the A-site or terminally stalled by mRNA-protein cross-links^25–27^ (Fig. S1C,D). Consistent with the above *in vivo* results, β-adrenergic stimulation of SVC-derived brown and white adipocytes also enhanced the expression of *Znf598* and other RQC-related genes (Fig. 1F,G). ZNF598 protein, the rate-limiting enzyme for RQC activity^28^, was elevated in BAT in response to cold challenge (Fig. 1H; Fig. S1E). Moreover, the abundance of ZNF598 was robustly increased in BAT and inguinal WAT (iWAT) of mice reared on a high-fat diet (HFD) for a short or prolonged time (Fig. 1I; Fig. S1F-H).

To monitor RQC engagement in adipocytes, we stably transfected T37i cells with a stalling reporter. The expressed mRNA accrues queues of ribosomes on a string of 20 AAA-based lysine codons that separates open reading frames (ORFs) for a green and a red fluorescent protein (GFP-20K-mKate2) (modified from^29^). The underlying principle of this assay is that mKate2 synthesis (mKate2/GFP ratio) is suppressed by ZNF598-mediated removal of queuing ribosomes on the stalling sequence. Defects in RQC activity or over-engagement of the pathway due to general collision stress at endogenous transcripts can be read out as an increase in the mKate2/GFP ratio. Stimulation of differentiated T37i BAT-like cells with CL and isoproterenol (Iso) increased mKate2-to-GFP ratio (Fig. 1J,K), indicating the appearance of ZNF598-engaging ribosome collisions on endogenous transcripts upon β-adrenergic stimulation. We also examined RQC activity with a modified reporter targeting the GFP-20K-mKate2 fusion protein to mitochondria (MitoGFP-20K-mKate2), which returned a similar result (Fig. 1L). These data suggest that β-adrenergic stimulation of adipocytes engages the RQC machinery to levels that impair dissociation of ribosomes from reporter-based stall sequences, despite elevated expression of RQC components. We next employed CRISPR technology to generate T37i cells KO (knockout) for *Znf598 (Znf598^-/-^)* (Fig. S1I). In differentiated T37i, proteomic profiling of ubiquitination site abundance revealed, among others, CL-induced and ZNF598-dependent ubiquitination of relevant lysines on RPS10 (Fig. 1M; Table S1). ZNF598-dependent RPS10 ubiquitination was also induced in differentiated T37i and differentiated 3T3L1 (WAT-like adipocyte precursor) upon stimulation with β-agonist, this time determined by western blot using the ribosome inhibitor anisomycin as a positive control (Fig. 1N,O, and Fig. S1J). Of note, differentiated but unstimulated T37i, unlike 3T3L1, displayed relatively high levels of RPS10 ubiquitination when assayed by this approach. Our results indicate that ribosome collision occurs upon thermogenic stimulation in adipocytes and that upregulation of the RQC pathway is evoked as a programmed protective response for cell-adaptive purposes.

### RQC is required for thermogenesis-associated enhanced translation of mitochondrial proteins and UCP1

To investigate whether ZNF598 affects the β-adrenergic stimulation-induced translational upregulation of mitochondrial genes, we analyzed total proteome changes in differentiated WT and *Znf598^-/-^* T37i cells. Principal component analysis (PCA) hinted that the large-scale proteome reprogramming induced by CL stimulation in these cells (separation according to principal component (PC) 1) were largely ZNF598-dependent (Fig. 2A). Comparing only CL-stimulated conditions, virtually all mitochondrial proteins as wells as UCP1 were less abundant in *Znf598^-/-^* cells compared to WT, suggesting that these cells are deficient for building mitochondrial mass (Fig. 2B; Table S2). Functional enrichment of protein networks also revealed that ZNF598 loss primarily negatively affected the abundance of mitochondrial proteins (Fig. 2C). Comparison of the mock-treated conditions revealed that these ZNF598-dependent differences could even be observed in resting T37i cells (Fig. S2A,B). Western blot analysis confirmed a clear ZNF598-dependent reduction in mitochondrial respiratory chain complexes and UCP1 in these cells, both under basal, CL- and Iso-stimulated conditions (Fig. 2D,E). We also studied SVC cells derived from adipose tissue stromal vascular fraction of mice with tamoxifen-inducible *Znf598* gene deletion (“*Znf598* cKO”, genotype *Rosa26-CreERT2 ; Znf598^fl/fl^*). When treated with tamoxifen and differentiated into brown adipocytes (Fig. 2F), these cells displayed modestly reduced abundance of the thermogenesis-related factors UCP1, DIO2 (Deiodinase type 2), and some respiratory chain proteins despite an incomplete targeting of the *Znf598* locus (Fig. 2G). Similar effect sizes were observed upon acute and incomplete knockdown of ZNF598 in T37i cells by siRNA (Fig. 2H). These differences in protein abundance were not mirrored by lower levels of mRNAs encoding mitochondrial proteins and thermogenesis-related factors. Rather, these mRNAs were enhanced in SVC-derived ZNF598-deficient brown adipocytes (Fig. S2C). In addition to their function in thermogenesis, white, beige, and brown adipocytes serve as dynamic energy repositories for the organism. Feeding, fasting, and energy demand in the periphery define the rate of lipid synthesis and degradation. Deletion of *Znf598* in T37i and differentiation into the BAT-like state did not affect the abundance of major proteins involved in lipogenesis (ACC and FASN), triglyceride synthesis (DGAT), fatty acid transport (CD36), or lipolysis (ATGL, HSL, and pHSL) under basal or β-adrenergic stimulated conditions (Fig. S2D). These results indicate that ZNF598 specifically regulates the translation of thermogenesis-related proteins without affecting unrelated aspects of adipocyte biology.

**Figure 2.**
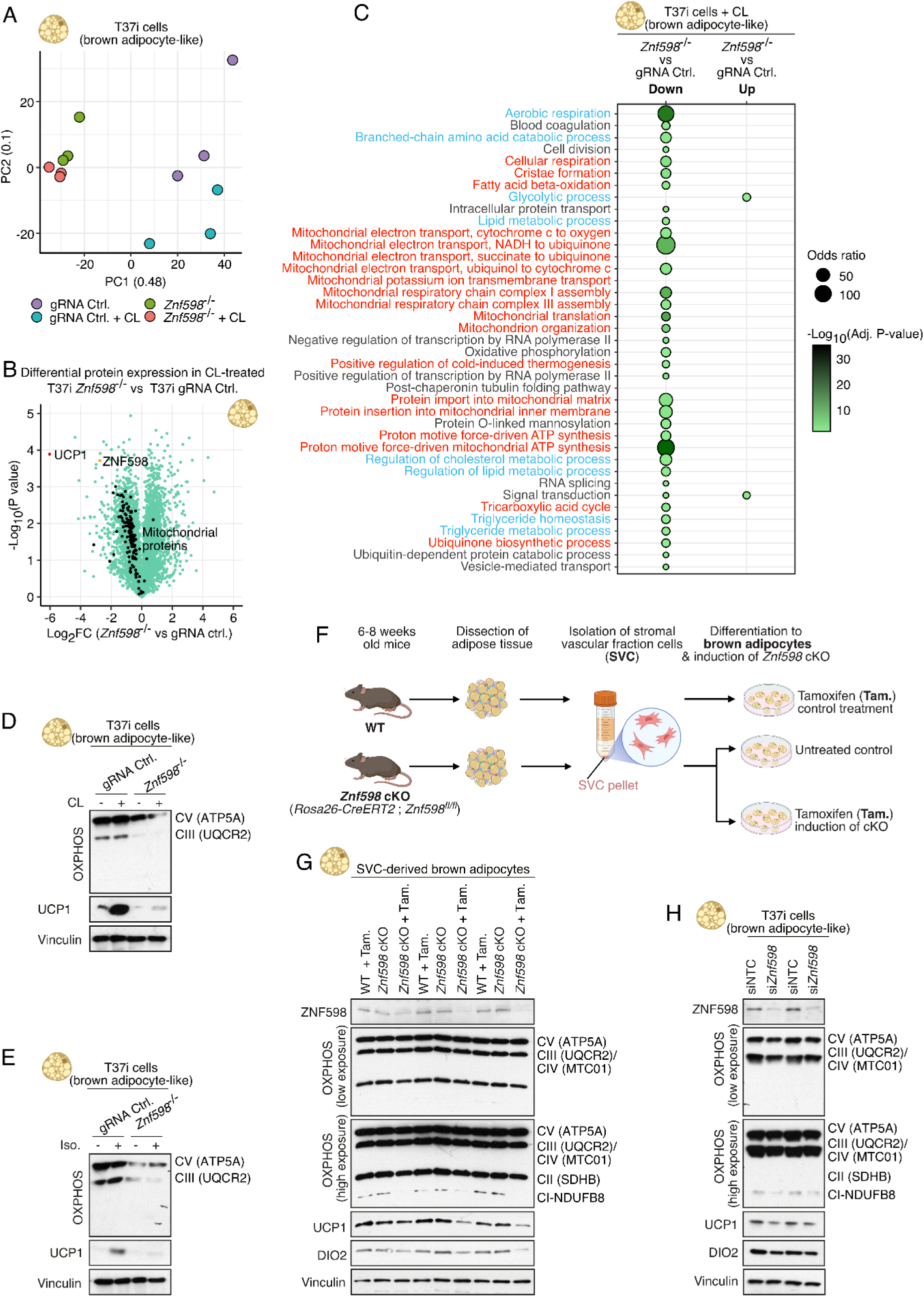
ZNF598 is essential for thermogenic programming and mitochondrial biogenesis in adipocytes. **(A)** Principal component analysis (PCA) of whole proteome quantification datasets from differentiated WT (gRNA Ctrl.) and *Znf598*^-/-^ T37i cells treated with or without β-agonist CL-316,243 (CL) treatment for 8 hours (n = 3 biological replicates). **(B)** Volcano plot comparing levels of individual proteins in WT and *Znf598*^-/-^ T37i brown adipocytes treated with CL (8 hours). ZNF598 (orange), UCP1 (red), and mitochondrial proteins (black) are highlighted. **(C)** Kyoto Encyclopedia of Genes and Genomes (KEGG) enrichment analysis of pathway differences based on whole proteome data from (B). Mitochondria-associated processes (red) and metabolic processes (blue) are highlighted. **(D–E)** Western blot (WB) analysis of protein lysates from WT and *Znf598*^-/-^ T37i brown adipocytes treated with CL **(D)** or isoproterenol (Iso.) **(E)** for 24 hours, using antibodies against OXPHOS, UCP1, and Vinculin. **(F)** Isolation of stromal vascular cells (SVC) and *ex vivo* generation of primary brown adipocytes deleted for *Znf598*. **(G–H)** WB analysis of protein lysates from **(G)** brown adipocytes derived from stromal vascular cells (SVC) isolated from WT and *Znf598* cKO mice (± Tamoxifen (Tam.) induction) and **(H)** T37i brown adipocytes depleted of ZNF598 by siRNA (si*Znf598*), using the indicated antibodies.

### ZNF598 is required for mitochondrial biogenesis and function in brown and beige adipocytes

All of the above indicate that RQC is required for mitochondrial function in adipocytes with thermogenic capacity. Indeed, ZNF598-deficient brown adipocytes were impaired in both basal respiration and maximal mitochondrial capacity, regardless of whether they originated from *Znf598* KO or *Znf598* siRNA-treated T37i cells or from SVC-derived ZNF598-deficient precursor cells, as measured by oxygen consumption rate by Seahorse technology (Fig. 3A,B; Fig. S3A). Moreover, T37i-derived brown adipocytes deleted for *Znf598* were refractory to β-agonist-induced boosting of the same parameters (Fig. 3C). Even 3T3L1-derived white adipocytes depleted for ZNF598 were refractory to the marginal boosting effects of β-agonist on mitochondrial function and abundance, but these cells did not display any differences under unstimulated conditions (Fig. S3B,C). All of these prompted us to investigate the impact of ZNF598 on mitochondrial acquisition upon β-adrenergic challenge. In SVC-derived brown adipocytes, deletion of *Znf598* blunted CL-induced increase in mitochondrial content, as indicated by mitotracker staining (Fig. 3D,E). siRNA-mediated depletion of ZNF598 in T37i-derived brown adipocytes resulted in a similar defect after challenge with two independent β-agonists, as indicated by both mitotracker staining and ratios of mitochondrial to genomic DNA (Fig. 3F-H).

**Figure 3.**
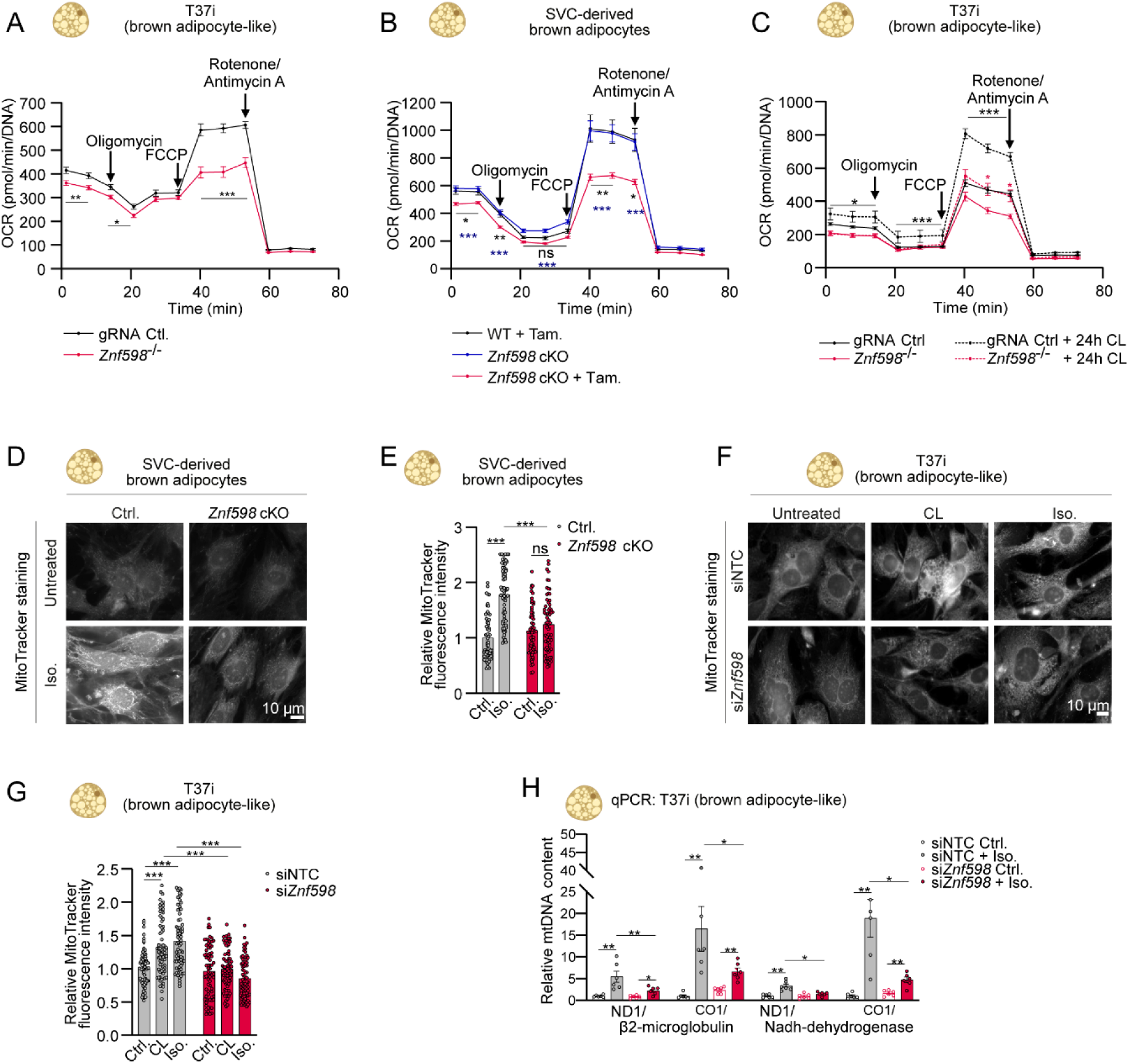
ZNF598 is required for β-adrenergic–stimulated mitochondrial respiration. **(A-C)** Seahorse analysis of mitochondrial respiration in **(A)** mock-treated WT (gRNA Ctrl.) and *Znf598*^-/-^ T37i brown adipocytes, **(B)** brown adipocytes from stromal vascular cells (SVC) isolated from WT and *Znf598* cKO mice (± Tamoxifen (Tam.) induction), and **(C)** WT and *Znf598*^-/-^ T37i brown adipocytes non-treated (NT) or treated with CL-316,243 (CL) for 24 hours. **(D–E)** Staining **(D)** and quantification of mitochondrial content by mitotracker fluorescence **(A)** in brown adipocytes derived from SVC isolated from WT and *Znf598* cKO mice (+ Tamoxifen induction) and treated with isoproterenol (Iso.) for 24 hours. Each data point represents the relative fluorescence intensity of an image field containing 10–15 cells. **(F–G)** As in (D) and (E), except that T37i-derived brown adipocytes were transfected with the indicated siRNAs (NTC, non-targeting control) and treated with CL and Iso. for 24 hours. **(H)** Quantitative PCR analysis of mitochondrial (mt)DNA levels relative to genomic DNA levels in T37i brown adipocytes depleted of ZNF598 by siRNA and treated with Iso. for 24 hours. Four independent primer sets targeting mtDNA were used. (A, B, C) All data points are an average of 11-19 replicates and (H) n = 6, corresponding to a technical duplicate of 3 biological replicates each. Data are plotted as mean, and all error bars represent the SEM. ns., non-significant; *, p≤0.05; **, p≤0.01; ***, p≤0.001 in Mann-Whitney test.

Brown adipocytes display higher mitochondrial content and respiration rate compared to white adipocytes. Of note, we detected the presence of ubiquitinated species of ZNF598 and RPS10 in SVC-derived brown (but not white) adipocytes (Fig. S3D,E), indicating that the RQC pathway is active in these cells. Consistently, unstimulated SVC-derived white adipocytes deleted for *Znf598* were not negatively affected with regard to mitochondrial respiration and abundance of mitochondrial complexes (Fig. S3F,G). Taken together, ZNF598 promotes mitochondrial biogenesis and function in brown, but not white, adipocytes at basal conditions. However, the RQC machinery is required for β-adrenergic-induced mitochondrial acquisition in both.

### Overexpression of ZNF598 boosts mitochondrial content, function, and uncoupling activity

ZNF598 was recently demonstrated to be a rate-limiting factor for RQC activity ^28^. Overexpression of ZNF598 in T37i-derived brown adipocytes was sufficient to elevate mitochondrial to genomic DNA ratio (Fig. S4A) and increase basal respiration, uncoupling activity, and maximal respiratory capacity of mitochondria (Fig. 4A). Electron microscopy-based analyses of the ultrastructure of these cells revealed an increased number and reduced size of mitochondria, especially those in contact with lipid droplets (Fig. 4B,C; Fig. S4B). These effects were accompanied by increased levels of respiratory complexes and elevated levels of UCP1 in ZNF598-overexpressing cells (Fig. 4D). As a likely consequence of the higher numbers of mitochondria, these cells also displayed lower levels of triglycerides (Fig. S4C,D). We also generated SVC-derived brown adipocytes with Tamoxifen-induced overexpression of ZNF598 (mouse genotype *Adipoq^CreERT^*^2^*; Rosa26^LSL-Flag-Znf5^*^98^, alias *Znf598^Adipoq-OE^*). When induced by tamoxifen prior to differentiation (Fig. 4E), these cells displayed elevated basal respiration, increased uncoupling activity, maximal respiratory capacity, increased abundance of UCP1 and respiratory complexes, and elevated mitochondrial to genomic DNA ratio and mitotracker signals (Fig. 4F,G; Fig. S4E), all indicators of increased mitochondrial mass and thermogenic activity. The effects of ZNF598 overexpression in T37i brown adipocytes were not further enhanced by β-adrenergic stimulation (Fig. 4H-J) and were not associated with changes in abundance of proteins involved in lipogenesis, triglyceride synthesis, lipid transport, and lipolysis (Fig. S4F). ZNF598 overexpression in SVC-derived brown adipocytes was also not associated with major changes in expression of the underlying genes (Fig. S4G). Finally, when overexpressed in SVC-derived white adipocytes, ZNF598 also had mild positive effects on basal respiration, uncoupling activity, and mitochondrial content (Fig. S4H-J) independent of effects on gene expression (Fig. S4K). Altogether, our results indicate that RQC activity can be boosted by overexpression of ZNF598 in brown adipocytes with positive effects on mitochondrial content, respiratory capacity, and uncoupling activity even in the absence of hormonal stimulation. This concept is on par with ZNF598, recently being described as a rate-limiting factor for RQC activity^28^.

**Figure 4.**
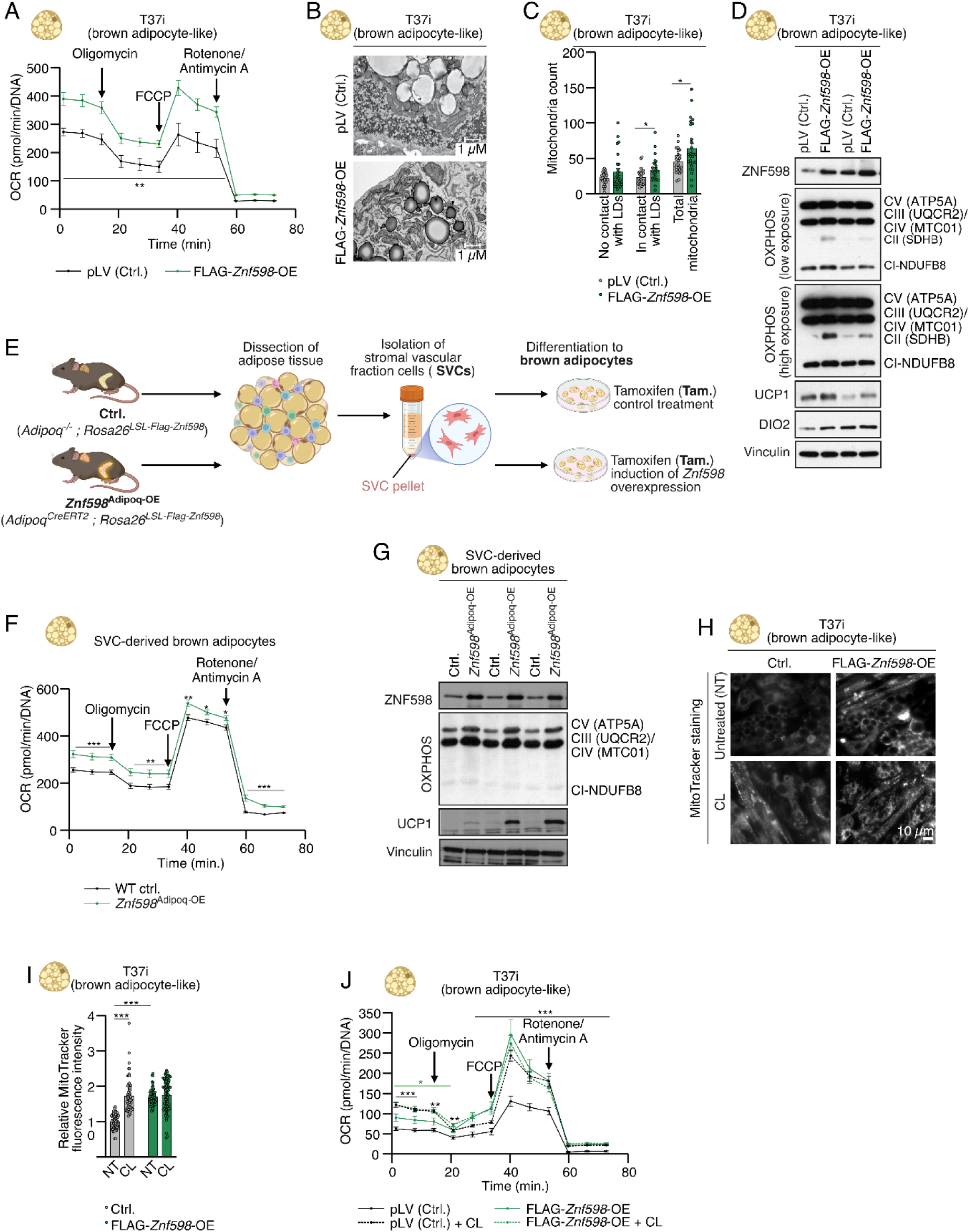
Overexpression of ZNF598 elevates respiration and mitochondrial content. **(A)** Seahorse analysis of mitochondrial respiration in T37i brown adipocytes stably transfected with empty vector (pLV) or overexpressing ZNF598. **(B-C)** Representative electron microscopy images **(B)** and quantification of the number of mitochondria per cell **(C)** of cells from (A). LD, Lipid droplets. **(D)** Western blot (WB) analysis of protein lysates from cells from (A), using antibodies against OXPHOS, UCP1, DIO2, and Vinculin. **(E)** *Ex vivo* generation of primary brown adipocytes conditionally overexpressing FLAG-ZNF598. Stromal vascular cells (SVC) were isolated from white adipose tissue (WAT) from mice with the indicated genotypes, treated with tamoxifen, and differentiated *in vitro* into brown adipocytes. **(F)** Seahorse analysis of mitochondrial respiration in brown adipocytes derived from SVC isolated from control mice and mice with tamoxifen-inducible overexpression of ZNF598 in adipocytes (*Znf598*^Adipoq-OE^). **(G)** Western blot (WB) analysis of protein lysates from cells from (E) and (F), using the indicated antibodies. **(H–I)** Staining **(H)** and quantification of mitochondrial content by mitotracker fluorescence **(I)** in cells from (A) non-treated (NT) or treated with CL-316,243 (CL) for 24 hours. Each data point represents the relative fluorescence intensity of an image field containing 10–15 cells. **(J)** Seahorse analysis of mitochondrial respiration in cells from (A) non-treated (NT) or treated with CL for 24 hours. (A, F, J) All data points are an average of more than 7 replicates. Data are plotted as mean, and all error bars represent the SEM. ns., non-significant; *, p≤0.05; **, p≤0.01; ***, p≤0.001 in Mann-Whitney test.

### Knockout of *Znf598* in adipocytes suppresses inducible iWAT browning and promotes glucose intolerance in mice

To study the effects of RQC in adipocytes *in vivo*, we generated mice carrying a conditional allele of *Znf598* (*Znf598*-flox). Excision of LoxP sites in *Znf598*-flox crossed with adipocyte-specific CRE-expressing mice (*Adipoq-cre*) resulted in the deletion of exons 2 to 5 (*Znf598^adipoq-cKO^*) (Fig. S5A) and a strong decrease of *Znf598* transcript levels in white and brown adipose tissues, but not in unrelated organs such as liver (Fig. S5B). These mice appeared phenotypically normal and were indistinguishable from their WT littermates with respect to weight and adipose tissue morphology. In order to gauge the plasticity of adipocytes *in vivo*, we injected mice with β-agonist CL over 10 days and examined iWAT for inducible browning (Fig. 5A). This response, determined as the appearance of multilocular adipocytes, was markedly suppressed in *Znf598^adipoq-cKO^* mice (Fig. 5B,C). On the other hand, BAT tissue appeared morphologically similar between genotypes (Fig. S5C). The apparent deficiency in the CL-induced browning response was accompanied by a similar negative effect on UCP1 upregulation and a reduction in the abundance of respiratory chain complexes, in iWAT but not BAT (Fig. 5D-G; Fig. S5D-F). Consistent with our *in vitro* data, these effects were not mirrored by differences in *Ucp1* mRNA levels (Fig. S5G,H), suggesting that CL-induced iWAT browning requires translation surveillance by the RQC pathway. The effects were not attributable to differences in whole-body and adipose tissue weights between genotypes in the experiment (Fig. S5I-L).

**Figure 5.**
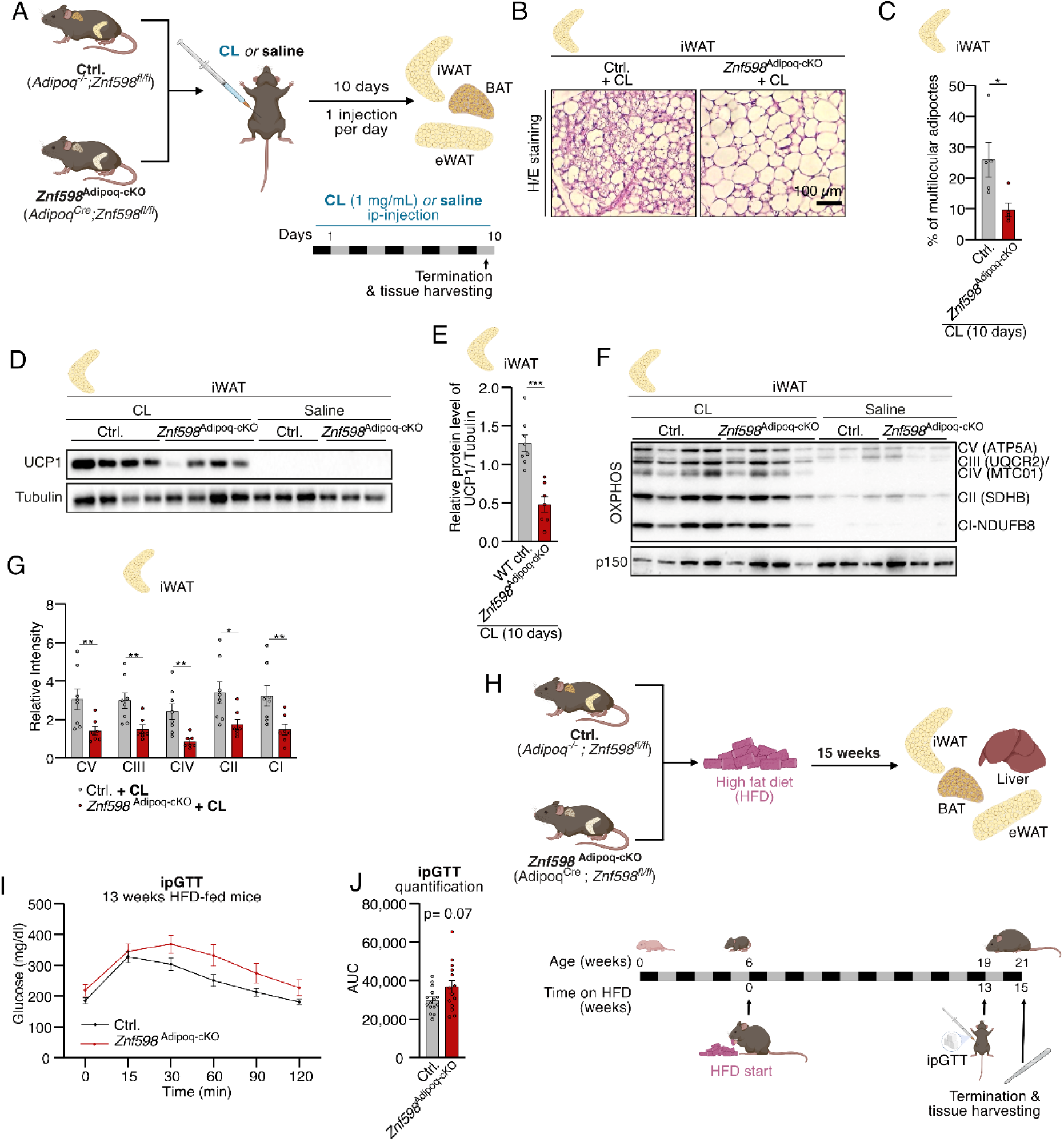
Adipocyte-specific deletion of ZNF598 impairs β-adrenergic stimulation-induced iWAT browning and promotes glucose intolerance in mice. **(A)** *In vivo* experimental strategy to study thermogenic responses. WT and *Znf598*^Adipoq-cKO^ (n = 8 and n = 7 biological replicates, respectively) mice were injected with CL-316,243 (CL) for 10 days before euthanasia. **(B)** Representative H/E (hematoxylin and eosin) staining of inguinal white adipose tissue (iWAT) in mice from (A). **(C)** Percentage of multilocular (brown/beige) adipocytes in iWAT of mice from (A). 5 mice from each group were randomly selected. **(D-E)** Western blot (WB) analysis **(D)** and corresponding densitometric quantification **(E)** of UCP1 protein levels in lysates from iWAT of mice from (A). **(F-G)** Western blot (WB) analysis **(F)** and corresponding densitometric quantification **(G)** of mitochondrial protein complexes in lysates from iWAT of mice from (A). **(H)** Schematic of high-fat diet (HFD)-feeding experiment. Mice with the indicated genotypes were changed from chow to HFD at 6 weeks of age. At 19 weeks of age, mice were challenged by Glucose tolerance test (ipGTT) and euthanized two weeks later. **(I)** ipGTT of mice with genotypes from (A) fed a high-fat diet (HFD) as in (H) (n = 15 and n = 14 biological replicates of Ctrl. and *Znf598*^Adipoq-cKO^, respectively). **(J)** Quantification of the area under the curve (AUC) for all mice in (I). Data are plotted as mean, and all error bars represent the SEM. ns., non-significant; *, p≤0.05; **, p≤0.01; ***, p≤0.001 in Mann-Whitney test.

Reduction in mitochondrial content in adipose tissue and diminished UCP1 abundance can predispose to metabolic disorders, including diabetes^2^. To evaluate whether *Znf598^adipoq-cKO^*mice were metabolically compromised in other aspects, we subjected mice to a high-fat diet (HFD), which induces obesity and glucose intolerance (Fig. 5H). Body weight gain, food intake, and energy expenditure were similar between genotypes during the feeding regimen (Fig. S5M-O), and we also did not observe any differences in organ weights and appearances at termination. However, when subjected to a glucose tolerance test (GTT), HFD-fed *Znf598^adipoq-cKO^* mice displayed both elevated basal blood glucose levels and defective glucose clearance (Fig. 5I,J). While not impacting on the development or morphological appearance of adipose tissue, our data indicate that *Znf598* deletion and loss of RQC activity in adipocytes compromise metabolic transitions in mice, here exemplified by inducible iWAT browning and blood glucose control.

### A mouse model with elevated ZNF598 expression in adipocytes displays increased thermogenic potential and improves glucose tolerance

To further investigate the influence on the RQC pathway on adipose tissue processes *in vivo*, we generated a mouse model with inducible and adipocyte-specific ZNF598 overexpression (*Znf598^Adipoq-OE^*) (Fig. S6A). Administration of tamoxifen to these mice resulted in a marked elevation of ZNF598 levels in adipose tissue (Fig. S6B). Subsequent injection of these mice with CL for 10 consecutive days (Fig. 6A) resulted in the expected decrease in body weight for both ZNF598-overexpressing and control mice without affecting food intake (Fig. S6C,D). However, *Znf598^Adipoq-OE^*mice displayed elevated energy dissipation throughout the experiment, reaching significance at day 9 of the injections (Fig. 6B). Moreover, overexpression of ZNF598 caused significant remodeling of the adipose tissue, where *Znf598^Adipoq-OE^* mice presented elevated BAT mass and a lower mass of epididymal WAT (eWAT) (Fig. 6C-E).

**Figure 6.**
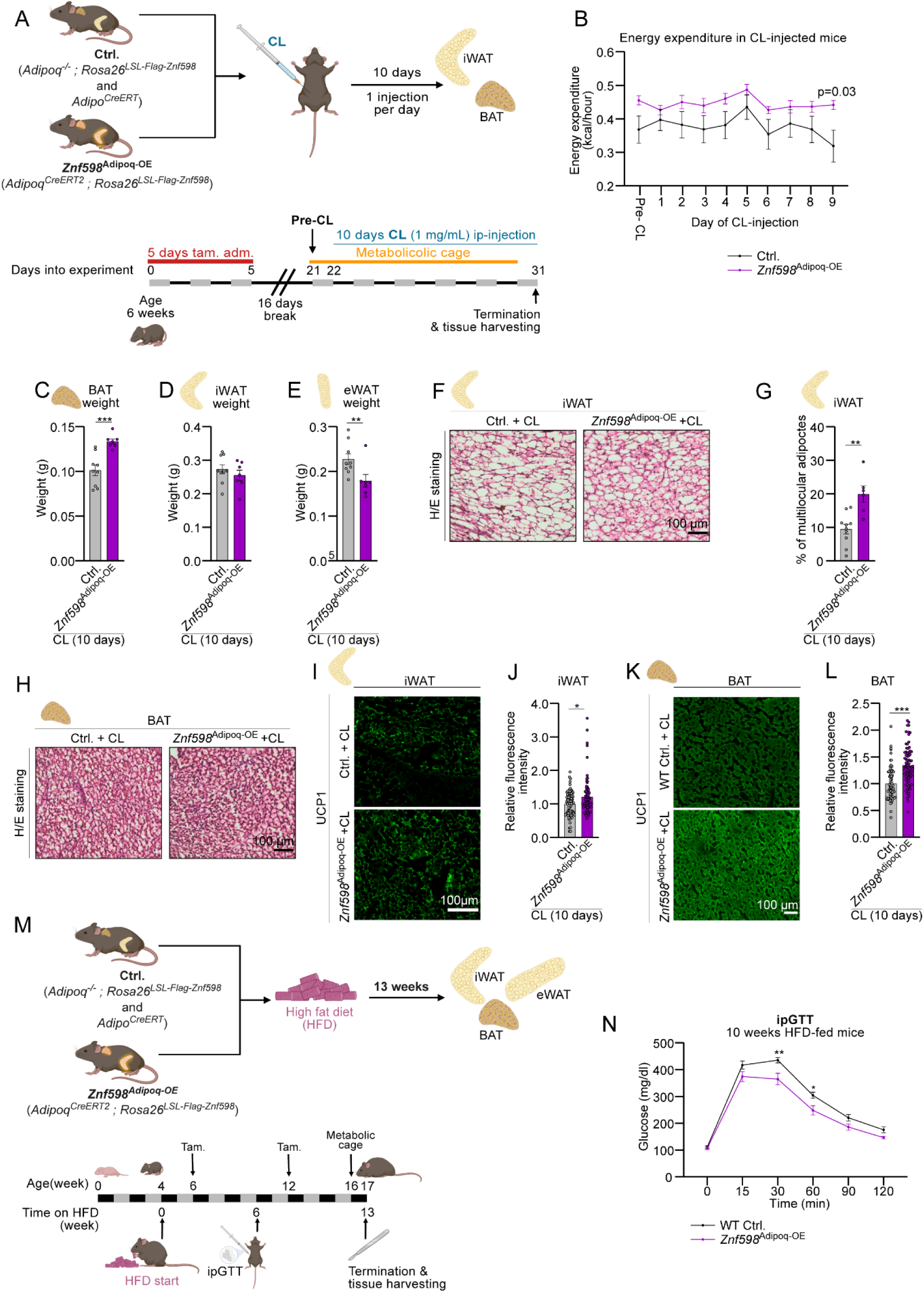
Tamoxifen-induced and adipocyte-specific overexpression enhances thermogenesis and protects from glucose intolerance in mice. **(A)** *In vivo* experimental strategy to study thermogenic responses in mice overexpressing ZNF598 in adipocytes. WT and *Znf598*^Adipoq-OE^ (n = 9 and n = 7 biological replicates, respectively) mice were injected with CL-316,243 (CL) for 10 days before euthanasia. **(B)** Whole-body energy expenditure of mice from (A) gavaged with tamoxifen or saline and injected as in (A). **(C-E)** Brown adipose tissue (BAT) **(C)**, inguinal white adipose tissue (iWAT) **(D)**, and epididymal WAT (eWAT) **(E)** weight at day 10 of the CL-injection experiment in (A). **(F)** Representative H/E (hematoxylin and eosin) staining of iWAT of mice from (A) **(G)** Percentage of multilocular (brown/beige) adipocytes in iWAT of mice from (A). **(H)** Representative H/E (hematoxylin and eosin) staining of BAT of mice from (A) **(I-J)** Representative immunostaining images **(I)** and quantification of UCP1 immunofluorescence signal per high-power field **(J)** of iWAT of mice from (A). **(K-L)** As in (I-J), except that BAT was immunostained **(K)** and quantified **(L)**. **(M)** Schematic of high-fat diet (HFD)-feeding experiment in mice overexpressing ZNF598 in adipocytes. Mice with the indicated genotypes (n = 15 and n = 8 biological replicates of Ctrl. and *Znf598*^Adipoq-OE,^ respectively) were changed from chow to HFD at 4 weeks of age and gavaged daily with tamoxifen during weeks 4 and 10. At 14 weeks of age, mice were challenged by Glucose tolerance test (ipGTT) and euthanized two weeks later. **(N)** ipGTT of mice with genotypes from (A) fed a high-fat diet (HFD) as in (M). Data are plotted as mean, and all error bars represent the SEM. ns., non-significant; *, p≤0.05; **, p≤0.01; ***, p≤0.001 in Mann-Whitney test.

These differences were associated with increased browning of iWAT in *Znf598^Adipoq-OE^* mice compared to controls, as determined by the number of multilocular adipocytes in this tissue (Fig. 6F,G). While BAT architecture was not influenced by *Znf598* overexpression (Fig. 6H), UCP1 abundance was elevated in both iWAT and BAT in these mice (Fig. 6I-L). These data indicate that elevation of RQC activity is sufficient to elevate energy expenditure in mice and to potentiate the impact of β-adrenergic stimulation on thermogenesis and iWAT browning.

We also subjected this mouse strain to an obesogenic diet and evaluated the metabolic consequences (Fig. 6M). Injection of tamoxifen initially after 2 weeks of HFD resulted in lower HFD-induced weight gain in ZNF598-overexpressing mice, but this difference compared to control animals disappeared over time (Fig. S6E). Mice were given a second boost of tamoxifen injection after 8 weeks and euthanized after 13 weeks of HFD feeding (Fig. 6M). While overexpression of ZNF598 in adipocytes did not significantly alter food intake, energy dissipation, or body composition (Fig. S6F-H), HFD-fed *Znf598^Adipoq-OE^*mice displayed improved blood glucose control after GTT challenge (Fig. 6N). Taken together, overexpression of ZNF598 in adipocytes protects against HFD-induced metabolic deterioration, resulting in an opposite phenotype to *Znf598^Adipoq-cKO^* mice.

## DISCUSSION

We demonstrate that increased translational demand during physiological tissue remodeling compromises translational fidelity and induces ribosomal collisions, thereby activating the RQC factor ZNF598, which is both necessary and sufficient to drive adaptive cellular remodeling. In particular, our results highlight that activation of the thermogenic program in adipocytes is associated with a remarkable induction of translational activity and ensuing ribosome collision. As a supportive part of this program, ZNF598 and a host of other RQC pathway components are induced to mitigate the potentially deleterious effects of ribosomal congestion on mitochondrial protein-encoding mRNAs and other relevant transcripts. Consequently, interference with the RQC pathway in several *in vitro* and *in vivo* models of adaptive thermogenesis is associated with defective cell and tissue responses. These include the acquisition of mitochondrial mass and induced iWAT browning.

Genetic defects in the RQC system are associated with neurodegenerative and neuronal disease in mice and humans^30^. Whether ribosome collision occurs on a large scale at basal conditions in other tissues or whether they manifest during physiologically relevant processes have not been appreciated. Here, we report for the first time both a physiological relevance of ribosome collision and a direct role of the RQC pathway in an extra-neuronal tissue. Our findings imply that activation of RQC is not only required for the induction of mitochondrial function and uncoupling activity in adipocytes but is also sufficient to drive these processes by selective stimulation of translation of mitochondrial proteins and UCP1. Our discoveries highlight RQC as a biological process that can be modulated to induce adaptive thermogenesis and combat obesity and associated diseases. A similar selective requirement for RQC was recently reported for human induced pluripotent stem cells (iPSCs). RQC is not required for maintenance of the pluripotent state but becomes critically important during differentiation^31^, which is likely associated with similar translation rewiring as in adipocytes undergoing thermogenic activation. Also, in intestinal stem cell differentiation, the metabolic shift from glycolysis to oxidative phosphorylation is brought about not by changes in gene expression but by rewiring of translational efficiency of the underlying transcripts^32^. This is also the case upon T cell activation, where the inverse metabolic shift is largely regulated at the translational rather than the transcriptional level ^33^. Whether these two cases of translationally regulated rapid changes to cellular identity are also associated with the induction of ribosome collision and a requirement for RQC remains to be studied.

Biochemical studies have indicated that the stoichiometry of RQC pathway components relative to translation initiation and elongation factors is highly unbalanced, with RQC components being significantly less abundant^34,35^. Our data highlight how β-adrenergic stimulation or dietary intervention increases the abundance of ZNF598 and other RQC components in adipocytes. However, as indicated by the higher rate of read-through of the stall sequence in an RQC reporter during β-adrenergic stimulation (Fig. 1J, L), the abundance of RQC components is not sufficient during the activation of the thermogenic activity of adipocytes. Previous studies indicated that engagement of the RQC pathway is critical in connection with mitochondrial stress ^29^. Our data from a range of *in vitro* and *in vivo* model systems highlights one physiological context where such perturbations occur and where RQC becomes critical. Deletion or silencing of *Znf598* resulted in defective upregulation of mitochondrial content and UCP1 abundance, without affecting the transcriptional induction of the underlying genes. Of note, mass spectrometry revealed UCP1 protein to be dramatically down-regulated in the absence of ZNF598 in T37i-derived brown adipocyte-like cells. While the reduced abundance of this protein might simply reflect the overall requirement of RQC for mitigating the damaging effects of rapid translational induction, it could also suggest the existence of a regulatory mechanism that directly affects the cellular production of UCP1 and mitochondrial proteins. ZNF598 is both sufficient and required for upregulation of UCP1 and mitochondrial proteins in both brown and white adipocytes.

In conclusion, our data highlight translational regulation as a key aspect of adaptive thermogenesis and demonstrate that ribosome collision is a bottleneck for the associated cellular remodeling. RQC and potentially other translation surveillance systems consequently become critical for translation of the key components determining the respiratory and thermogenic capacity of beige and brown adipocytes. These associations may have wider implications for physiologically relevant cellular transitions that require rapid remodeling of the proteome.

## ACKNOWLEDGEMENTS

We thank Drs. Malgorzata Alicja Sliwinska and Hanna Nieznańska from the Laboratory of Imaging Tissue Structure and Function for providing Transmission Electron Microscopy service, and Dr. Jedrzej Szymanski from the Laboratory of Imaging Tissue Structure and Function (both located at the Nencki Institute for Experimental Biology) for help with imaging of adipocytes. This study was supported by the Dioscuri Center of Scientific Excellence (grant number UMO-2018/01/H/NZ4/00002) – the programme initiated by the Max Planck Society (MPG), jointly managed with the National Science Center in Poland (NCN), and mutually funded by the Polish Ministry of Science and Higher Education (MNSW) and the German Federal Ministry of Research, Technology, and Space (BMFTR) granted to Grzegorz Sumara. Some of the initial results were obtained with the support of a Minatura pre-grant from the NCN (nr. 2022/06/X/NZ3/00539) awarded to Toufic Kassouf. Work in the Bekker-Jensen lab was supported by the European Research Council (ERC) under the European Union’s Horizon 2020 research and innovation program (grant agreement 863911 - PHYRIST) and Independent Research Fund Denmark (grant no. 3101-00344B). Center for Gene Expression (CGEN) is a Center of Excellence funded by The National Danish Research Foundation (grant no. DNRF166). Mass spectrometry-based proteomics analyses were performed by the Proteomics Research Infrastructure (PRI) at the University of Copenhagen (UCPH), supported by the Novo Nordisk Foundation (NNF) (grant agreement number NNF19SA0059305).

---

**Figure S1.**
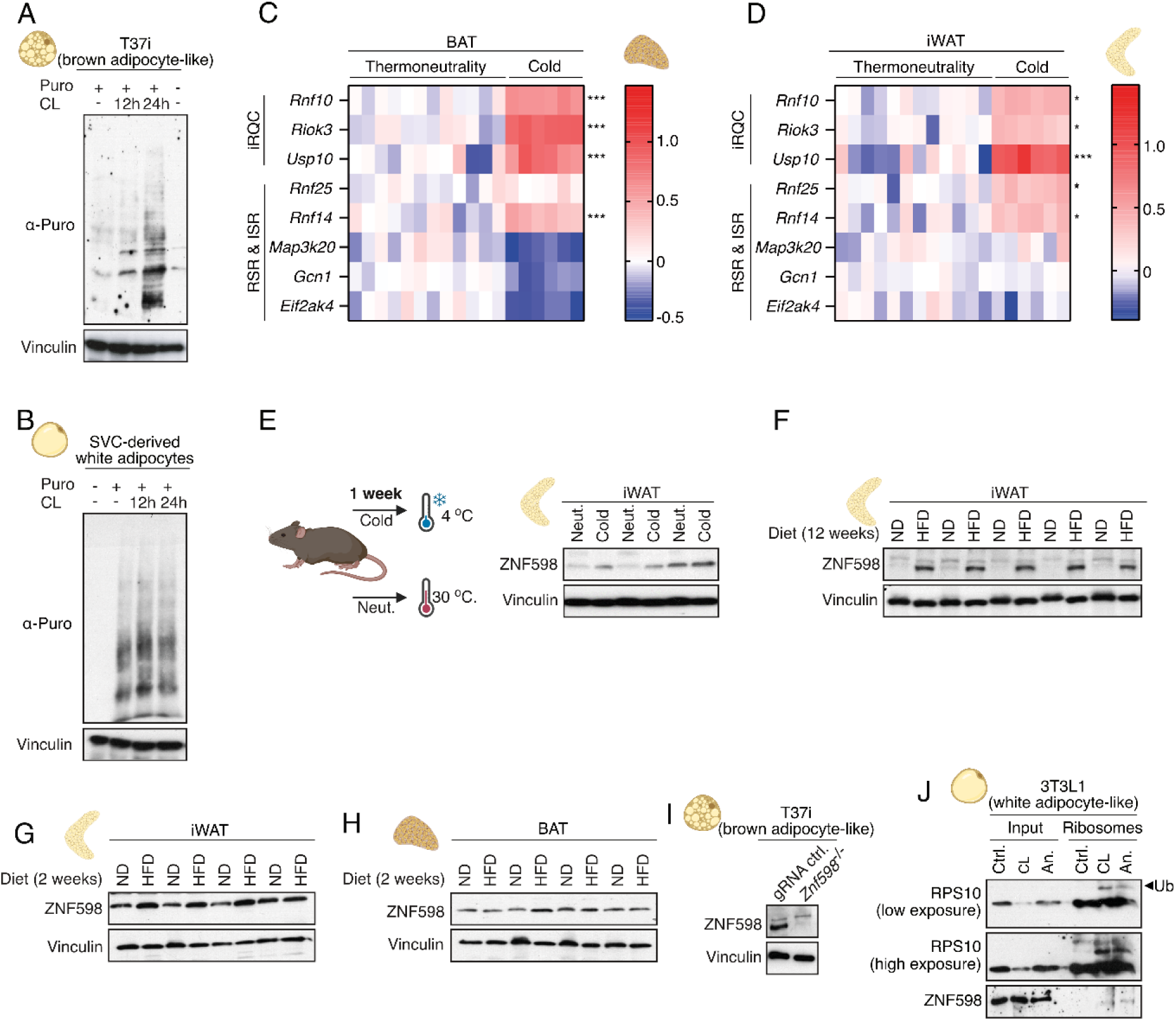
Translational Activation and Ribosome Collision in Thermogenic Adipocytes. **(A-B)** Determination of ribosomal output using puromycin-labeling of proteins in brown adipocytes derived from T37i cells **(A)** and white adipocytes derived from stromal vascular cells (SVC) **(B)** treated with CL-316,243 (CL) for the indicated time. **(C-D)** Heat maps showing the log2-transformed fold changes of mRNA levels for proteins functioning in initiation ribosome-associated quality control (iRQC), ribotoxic stress response (RSR) and integrated stress response (ISR) in brown adipose tissue (BAT) **(C)** and inguinal white adipose tissue (iWAT) **(D)** of mice kept at room temperature or exposed to 4 °C (n = 12 and n = 6 biological replicates, respectively) for 24 hours. **(E)** Western blot (WB) analysis of the abundance of ZNF598 in iWAT collected from WT mice following 1 week of thermoneutrality (30 °C, neut.) or cold exposure (4 °C, cold). **(F)** WB analysis of the abundance of ZNF598 in iWAT of WT mice fed a normal chow diet (ND) or a high-fat diet (HFD) for 12 weeks. **(G-H)** WB analysis of the abundance of ZNF598 in iWAT **(G)** and BAT **(H)** of mice fed ND or HFD for 2 weeks. **(I)** WB analysis of the abundance of ZNF598 in WT (gRNA Ctrl.) and *Znf598* knockout (^-/-^) T37i cells. **(J)** Western blot (WB) analysis of protein lysates (input) and pelleted ribosomes from 3T3L1 adipocytes treated with CL (24 hours) or anisomycin (An, 1 hour) as a positive control. (C, D) *, p≤0.05; ***, p≤0.001 in student’s t-test.

**Figure S2.**
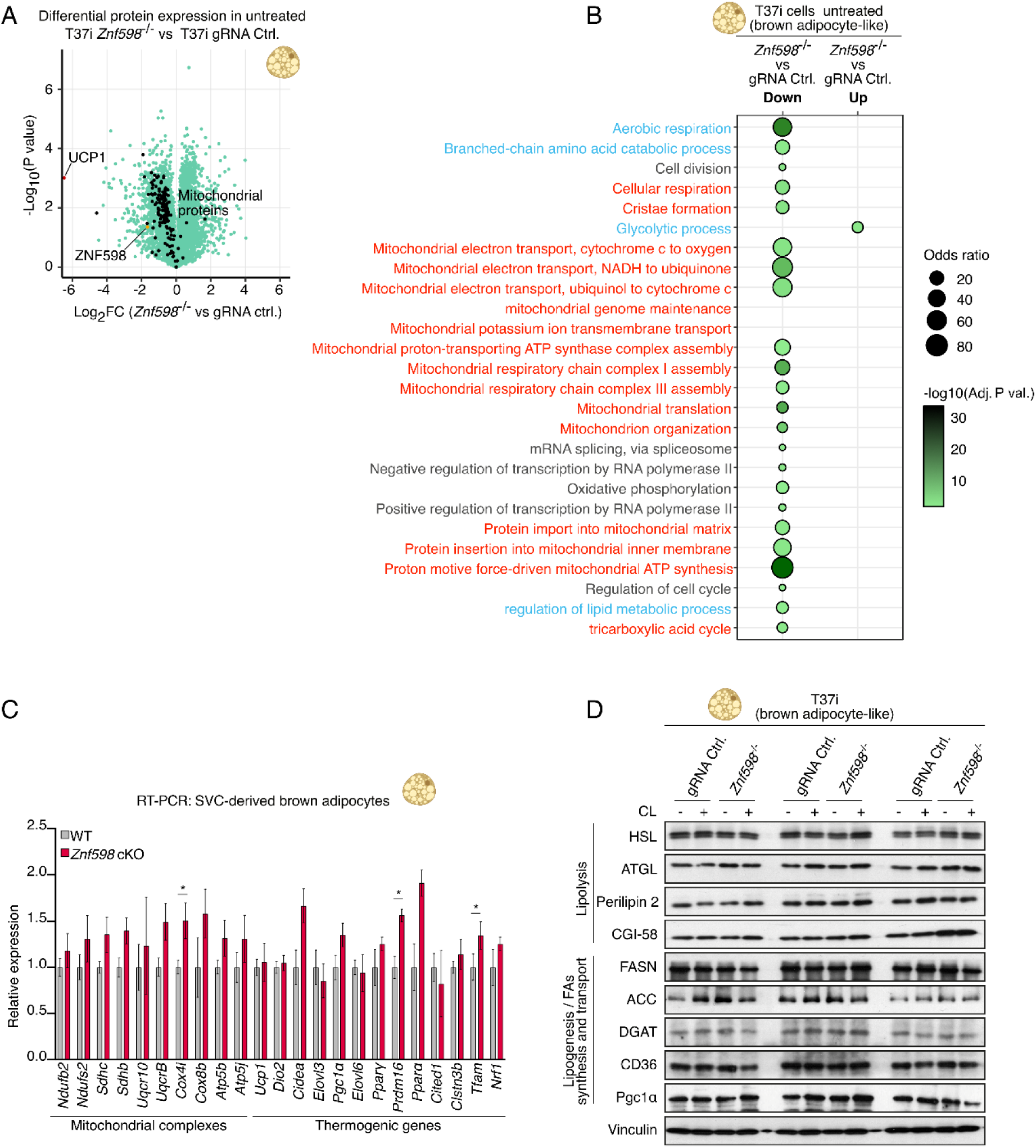
Reduced levels of mitochondrial and thermogenic proteins in ZNF598-deficient adipocytes. **(A)** Volcano plot comparing levels of individual proteins in mock-treated WT (gRNA Ctrl.) and *Znf598*^-/-^ T37i brown adipocytes. ZNF598 (orange), UCP1 (red), and mitochondrial proteins (black) are highlighted. **(B)** Kyoto Encyclopedia of Genes and Genomes (KEGG) enrichment analysis of pathway differences based on whole proteome data from (A). Mitochondria-associated processes (red) and metabolic processes (blue) are highlighted. **(C)** RT-PCR analysis for expression of mitochondrial and thermogenic genes in brown adipocytes derived from stromal vascular cells (SVC) isolated from WT and *Znf598* cKO mice (+ Tamoxifen induction). Relative expression values (Ctrl. = 1) are plotted as mean, and all error bars represent the SEM (n = 4 biological replicates). *, p≤0.05 in Mann-Whitney test. **(D)** Western blot (WB) analysis of protein lysates from WT and *Znf598*^-/-^ T37i brown adipocytes treated with CL-316,243 (CL) for 24 hours, using the indicated antibodies.

**Figure S3.**
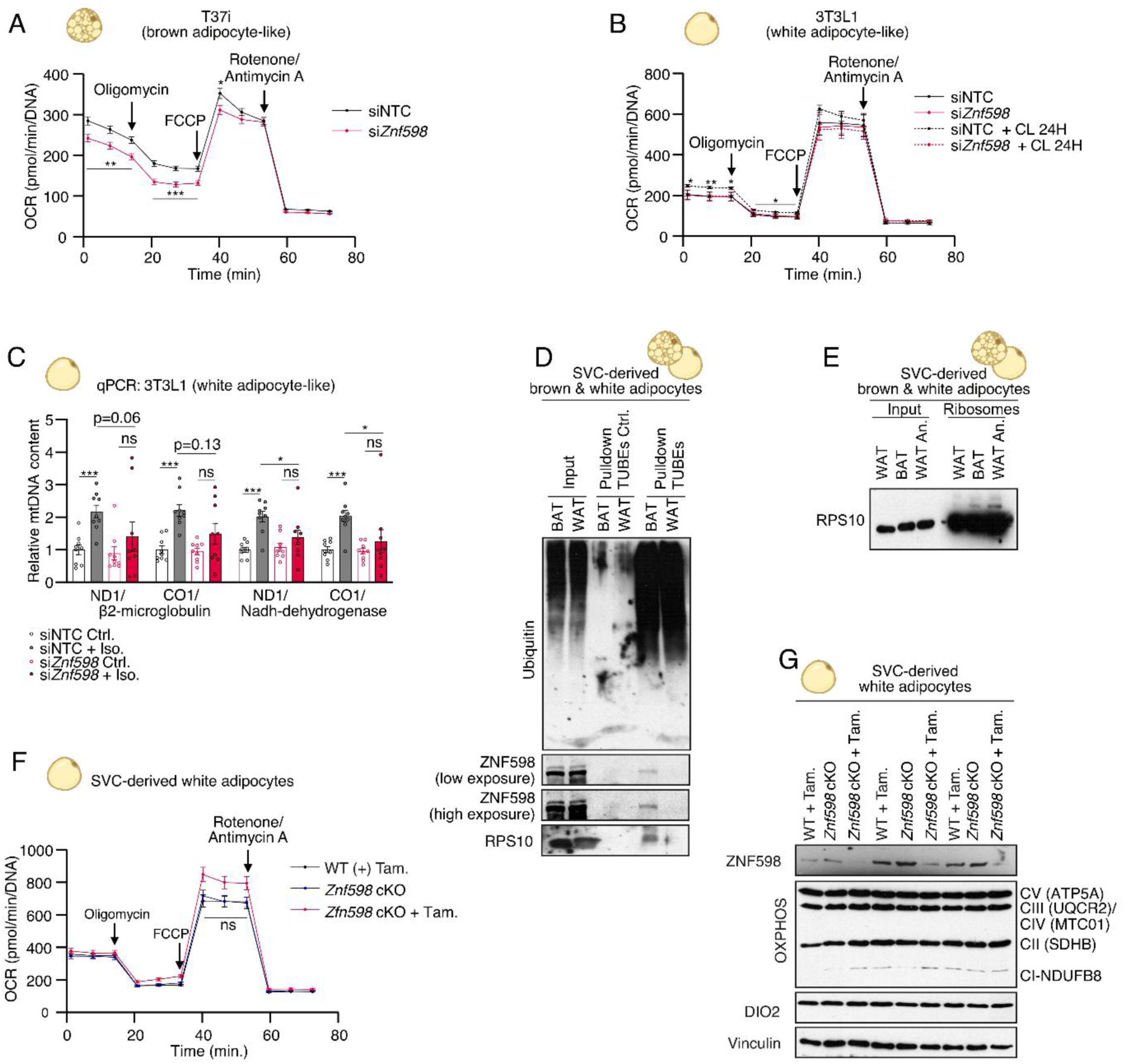
β-adrenergic stimulation marginally promotes ZNF598-dependent respiration in white adipocytes. **(A-B)** Seahorse analysis of mitochondrial respiration in **(A)** T37i brown adipocytes depleted of ZNF598 by siRNA (si*Znf598*) or transfected with non-targeting control (NTC) and **(B)** 3T3L1 white adipocytes depleted of ZNF598 by siRNA (si*Znf598*) or transfected with non-targeting control (NTC), either non-treated (NT) or treated with CL-316,243 (CL) for 24 hours. **(C)** Quantitative PCR analysis of mitochondrial (mt)DNA levels relative to genomic DNA levels in 3T3L1-derived white adipocytes depleted of ZNF598 by siRNA and treated with Iso. for 24 hours. Four independent primer sets targeting mtDNA were used. **(D)** Enrichment of ubiquitinated proteins by Agarose-TUBE2 pulldown from brown (BAT) and white (WAT) adipocytes derived from mouse stromal vascular cells (SVC). Western blot (WB) analysis of protein lysates (input) and pulldowns using antibodies against Ubiquitin, ZNF598, and RPS10. **(E)** WB analysis of protein lysates (input) and pelleted ribosomes from WAT and BAT from (D). WAT cells were treated with anisomycin (An., 1 hour) as a positive control. **(F)** Seahorse analysis of cellular respiration and **(G)** WB analysis of protein lysates from white adipocytes derived from SVC isolated from WT and *Znf598* cKO mice (± Tamoxifen (Tam.) induction). (A, B, F) All data points are an average of 12-22 replicates and (C) n = 9, corresponding to technical triplicates of 3 biological replicates. Data are plotted as mean, and all error bars represent the SEM. ns., non-significant; *, p≤0.05; ***, p≤0.001 in Mann-Whitney test.

**Figure S4.**
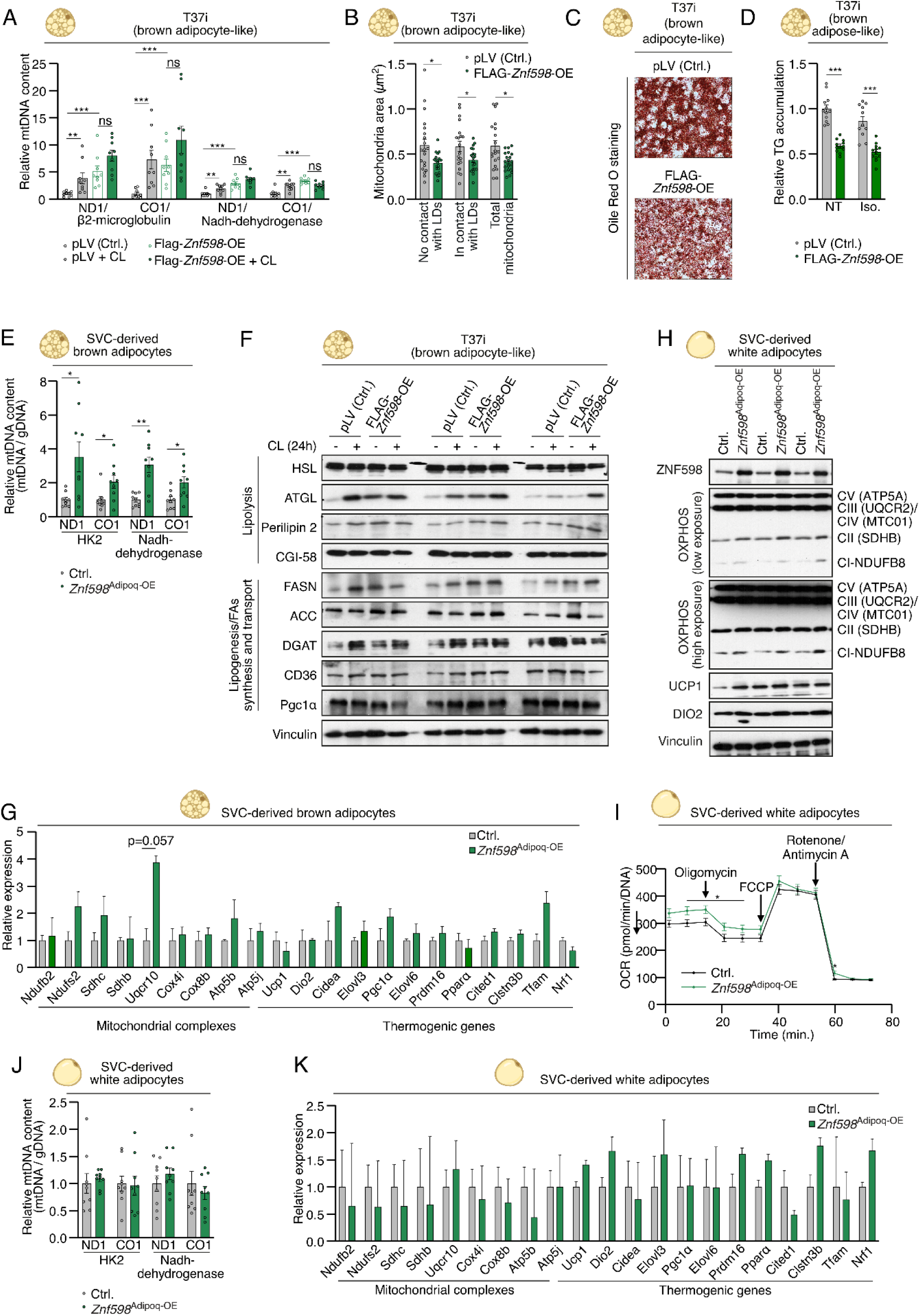
Effects of ZNF598 overexpression on levels of thermogenic and mitochondrial proteins. **(A)** Quantitative PCR analysis of mitochondrial (mt)DNA levels relative to genomic DNA levels in T37i brown adipocytes stably transfected with empty vector (pLV) or overexpressing ZNF598. Cells were left non-treated (Ctrl.) or treated with CL-316,243 (CL) for 24 hours. Four independent primer sets targeting mtDNA were used. Data are plotted as relative expression values (Non-treated Ctrl. = 1). **(B)** Quantification of mitochondrial area in electron microscopy images of cells from (A). LD, Lipid droplets. **(C-D)** Representative images of staining **(C)** and quantification **(D)** of cells from (A) for triglycerides and other neutral lipids using AdipoRed reagent. Data are plotted as relative values (Non-treated Ctrl. = 1). **(E)** As in (A), except for brown adipocytes derived from stromal vascular cells (SVC) isolated from control mice and mice with tamoxifen-inducible overexpression of ZNF598 in adipocytes (Znf598^Adipoq-OE^). **(F)** Western blot (WB) analysis of protein lysates from cells from (A). Cells were left non-treated or treated with CL for 24 hours. **(G)** RT-PCR analysis for expression of mitochondrial and thermogenic genes in brown adipocytes from (E). Data are plotted as relative expression values (Ctrl. = 1). **(H)** Western blot (WB) analysis of protein lysates from white adipocytes derived from SVC isolated from control mice and mice with tamoxifen-inducible overexpression of ZNF598 in adipocytes (*Znf598*^Adipoq-OE^), using antibodies against ZNF598, OXPHOS, UCP1, DIO2, and Vinculin. **(I)** Seahorse analysis of mitochondrial respiration in white adipocytes from (H). **(J)** As in (E), using cells from (H). **(K)** As in (G), using cells from (H). (A, E, J) n = 9 corresponding to technical triplicate of 3 biological replicates, (D) n = 11-12 from 3 biological replicates, (G, K) n = 3-4 biological replicates, and (I) all data points are an average of 23 (Ctrl.) or 36 (Znf598^Adipoq-OE^) replicates. Data are plotted as mean, and all error bars represent the SEM. ns., non-significant; *, p≤0.05; **, p≤0.01; ***, p≤0.001 in Mann-Whitney test.

**Figure S5.**
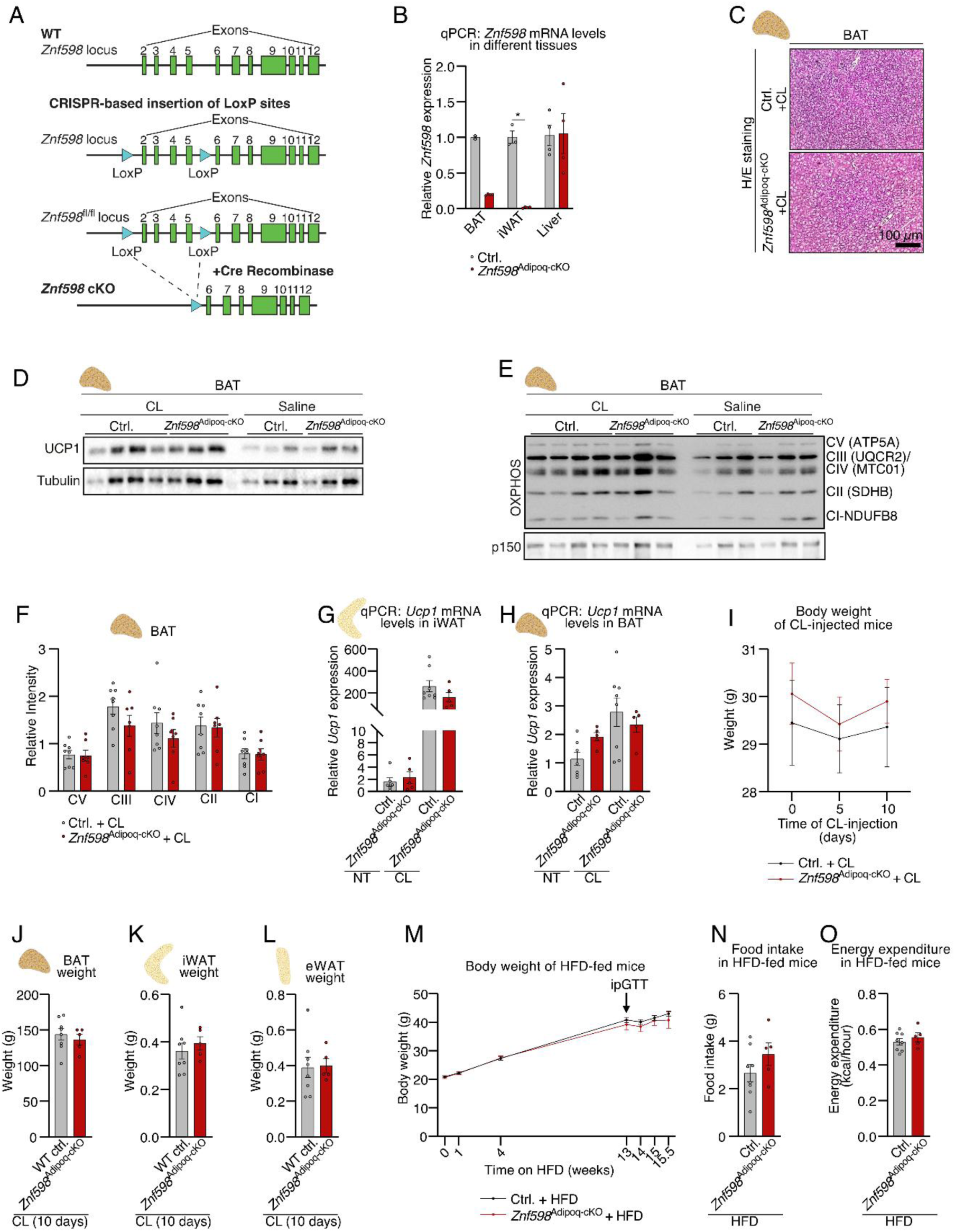
CL-316,243 injection does not affect energy balance in *Znf598*^Adipoq-cKO^ mice. **(A)** Schematic of the floxed *Znf598* allele and its targeting by CRE-LoxP in *Znf598*^Adipoq-cKO^ mice. **(B)** qPCR analysis of relative *Znf598* mRNA levels in brown adipose tissue (BAT), inguinal white adipose tissue (iWAT), and liver in WT and *Znf598*^Adipoq-KO^ mice. **(C)** Representative H/E (hematoxylin and eosin) staining of BAT from WT and *Znf598*^Adipoq-cKO^ mice after injection of CL-316,243 (CL) or saline for 10 days. **(D)** Western blot (WB) analysis of UCP1 protein levels in lysates from BAT of mice from (A) treated as in (C). **(E-F)** WB analysis **(E)** and corresponding densitometric quantification **(F)** of mitochondrial protein complexes in lysates from BAT of mice from (A) treated as in (C). **(G-H)** qPCR analysis of relative *Ucp1* mRNA levels in iWAT **(G)** and BAT **(H)** of mice from (A) treated as in (C). **(I)** Changes in body weight of mice from (A) over the course of the CL-injection experiment in (C). **(J-L)** BAT **(J)**, iWAT **(K)**, and eWAT (epididymal white adipose tissue) **(L)** weight at day 10 of the CL-injection experiment in (C). **(M-O)** Body weight evolution **(M)**, food intake **(N)**, and energy expenditure **(O)** of WT and *Znf598*^Adipoq-cKO^ mice fed a high-fat diet (HFD) for 15.5 weeks. (M) n = 15 and n = 14 biological replicates of Ctrl. and *Znf598*^Adipoq-cKO,^ respectively. (N-O) n = 8 Ctrl. and n = 5 biological replicates of Ctrl. and *Znf598*^Adipoq-cKO^, respectively, randomly selected from the main experimental group. Data are plotted as mean, and all error bars represent the SEM. ns., non-significant; *, p≤0.05; **, p≤0.01; ***, p≤0.001 in Mann-Whitney test.

**Figure S6.**
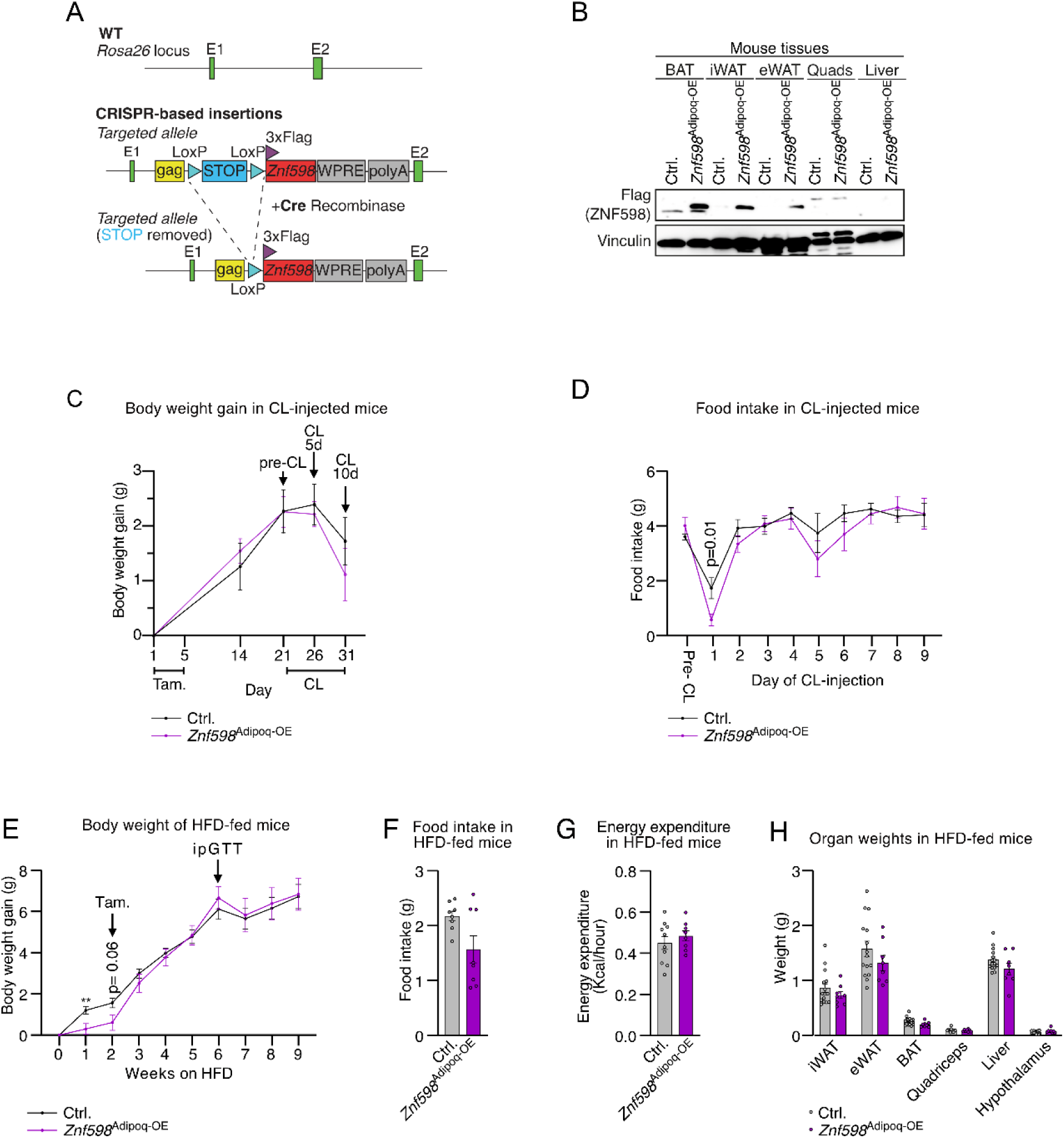
Overexpression of ZNF598 in adipose tissue does not affect body composition in HFD-fed mice. **(A)** Schematic of the Rosa26-integrated *FLAG-Znf598* transgene and its activation by CRE-LoxP upon crossing with AdiponectinCreERT in *Znf598*^Adipoq-OE^ mice. Recombination and transgene expression is activated by tamoxifen administration in this model. STOP, stop codon; WPRE, Woodchuck Hepatitis Virus Posttranscriptional Regulatory Element. **(B)** Western blot (WB) analysis of FLAG-ZNF598 expression in the indicated adipose tissues, quadriceps muscle (Quad.), and liver in mice from (A) gavaged with tamoxifen or control (Ctrl.) mice. **(C-D)** Changes in body weight **(C)** and food intake **(D)** of *Znf598*^Adipoq-OE^ mice injected with CL-316,243 (CL) (n = 15 and n = 8 biological replicates of Ctrl. and *Znf598*^Adipoq-OE^, respectively). **(E)** Change in bodyweight during the first 9 weeks of high-fat diet (HFD)-feeding of *Znf598*^Adipoq-OE^ mice. **(F-H)** food intake **(F)**, energy expenditure **(G)**, and weight of indicated organs **(H)** at termination of *Znf598*^Adipoq-OE^ mice fed HFD for 12 weeks (n = 8 biological replicates randomly selected from each experimental group). Data are plotted as mean, and all error bars represent the SEM. ns., non-significant; *, p≤0.05; **, p≤0.01; ***, p≤0.001 in Mann-Whitney test.

